# Complex roles for proliferating cell nuclear antigen in restricting human cytomegalovirus replication

**DOI:** 10.1101/2025.01.06.631530

**Authors:** Pierce Longmire, Olivia Daigle, Sebastian Zeltzer, Matias Lee, Marek Svoboda, Marco Padilla-Rodriguez, Carly Bobak, Giovanni Bosco, Felicia Goodrum

**Affiliations:** Graduate Program in Molecular Medicine, University of Arizona, Tucson, Arizona, USA; Department of Immunobiology, University of Arizona, Tucson, Arizona, USA; BIO5 Institute, University of Arizona, Tucson, Arizona, USA; Department of Molecular and Systems Biology, Dartmouth Geisel College of Medicine, Hanover, New Hampshire, USA; Research Computing and Data Services, Information, Technology, and Consulting, Dartmouth College, Hanover, New Hampshire, USA; Microscopy Shared Resource, University of Arizona Cancer Center, Tucson, Arizona, USA

**Author notes:** Address correspondence to Felicia Goodrum.

## Abstract

DNA viruses at once elicit and commandeer host pathways, including DNA repair pathways for virus replication. Despite encoding its own DNA polymerase and processivity factor, human cytomegalovirus (HCMV) recruits the cellular processivity factor, proliferating cell nuclear antigen (PCNA) and specialized host DNA polymerases involved in translesion synthesis (TLS) to replication compartments (RCs) where viral DNA (vDNA) is synthesized. While the recruitment of TLS polymerases is important for viral genome stability, the role of PCNA is poorly understood. PCNA function in DNA repair is regulated by monoubiquitination (mUb) or SUMOylation of PCNA at lysine 164 (K164). We find that mUb-PCNA increases over the course of infection, and modification of K164 is required for PCNA-mediated restriction of virus replication. mUb-PCNA plays important known roles in recruiting TLS polymerases to DNA, which we have shown are important for viral genome integrity and diversity, represented by novel junctions and single nucleotide variants (SNVs), respectively. We find that PCNA drives SNVs on vDNA similar to Y-family TLS polymerases, but that this did not require modification at K164. Unlike TLS polymerases, PCNA was dispensable for preventing large scale rearrangements on vDNA. These striking results suggest separable PCNA-dependent and - independent functions of TLS polymerases on vDNA. By extension, these results imply roles for TLS polymerase beyond their canonical function in TLS in host biology. These findings highlight PCNA as a complex restriction factor for HCMV infection, likely with multiple distinct roles, and provides new insights into the PCNA-mediated regulation of DNA synthesis and repair in viral infection.

**IMPORTANCE:** Genome synthesis is a critical step of virus life cycles and a major target of antiviral drugs. Human cytomegalovirus, like other herpesviruses, encodes machinery sufficient for viral DNA synthesis and relies on host factors for efficient replication. We have shown that host DNA repair factors play important roles in HCMV replication, but our understanding of this is incomplete. Building on previous findings that specialized host DNA polymerases contribute to HCMV genome integrity and diversity, we sought to determine the importance of PCNA, the central polymerase regulator. PCNA associates with nascent viral DNA and restricts HCMV replication. While PCNA is dispensable for genome integrity, it contributes to genome diversity. Our findings suggest that host polymerases function on viral genomes by separable PCNA-dependent and - independent mechanisms. Through revealing complex roles for PCNA in HCMV replication, this study expands the repertoire of host DNA synthesis and repair proteins hijacked by this ubiquitous herpesvirus.

## INTRODUCTION

Human cytomegalovirus (HCMV) is a ubiquitous herpesvirus that has co-evolved with humans and persists in a majority of the world’s population through establishment of latent infection. Latency is defined as a quiescent infection where viral genomes are maintained in the absence of viral genome synthesis and viral progeny production (1). For HCMV, progenitor cells of the myeloid lineage and monocytes have been described as a major reservoir for latency (2, 3). While most seropositive individuals experience asymptomatic infection, HCMV reactivation from latency poses serious disease and mortality risk for immunocompromised individuals including solid organ and stem cell transplant recipients. Infection during pregnancy and viral transmission to the fetus occurs in 1 in 200 births in the United States. Of these newborns with congenital infection, 20% will develop permanent disabilities, making HCMV the leading cause of infectious disease-related birth defects in the United States (4, 5). There are no clinical strategies to target or control latent HCMV infection due to limited knowledge of how the virus toggles between latent and replicative states. Complex interplay between viral determinants and host biology is central to regulation of viral gene expression, genome amplification, and decision to establish latency or replicate (6). Consequently, investigating these virus-host interactions at a molecular level is important to building a mechanistic understanding of latency and reactivation.

While HCMV encodes several proteins sufficient for viral DNA (vDNA) synthesis, it also relies on its host for specific factors that may fine-tune or modulate specific aspects of synthesis. HCMV infection drastically alters cell cycle progression (7) and induces E2F1 transcription factor activity, which regulates many host DNA synthesis and repair genes (8).

Virus replication also activates DNA damage response signaling through major kinases, ataxia-telangiectasia mutated (ATM) and ataxia-telangiectasia Rad3-related (ATR) (9). Many of the downstream repair proteins are re-localized to nuclear viral replication compartments (RCs), sites of vDNA synthesis, but their specific contributions to the viral replication cycle are poorly understood (8–11). In previous studies, we found that HCMV recruits specialized host DNA polymerases involved in translesion synthesis (TLS) pathways to modulate virus replication despite encoding its own DNA polymerase, *UL54*. TLS is a DNA damage bypass pathway in which specialized polymerases are recruited to replication forks or sites of damage in order to synthesize through lesions and prevent replication fork stalling or collapse (12). TLS polymerases lack proofreading activity and have a warped active site compared to replicative DNA polymerases, which allows for synthesis across DNA lesions albeit with lower fidelity (13). Therefore, TLS is a double-edged sword in which bulky nucleotide lesions may be bypassed during replication in a process that comes at the cost of potentially introducing small point mutations into newly synthesized DNA. We specifically found that the Y-family polymerases eta (ρι), kappa (κ), and iota (1) restrict vDNA synthesis and viral replication, while the B-family polymerase zeta (ζ) and its putative scaffold, Y family polymerase Rev1, are required for efficient vDNA synthesis and virus replication. Despite these polymerases having opposite effects on viral replication, all are required for viral genome integrity whereby the depletion of either group of polymerases results in a significant increase in aberrant recombination and novel DNA junctions, primarily inversions, across the viral genome (14).

Proliferating cell nuclear antigen (PCNA) is an essential sliding clamp that acts as a processivity factor, maintaining polymerase association with DNA and promoting processivity of DNA synthesis (15). Normally, PCNA interacts with B-family DNA polymerases delta (δ) and epsilon (ε) through a PCNA interacting protein (PIP) motif (15). At sites of damage, PCNA is monoubiquitinated (mUb-PCNA) at lysine reside 164 (K164) by the E3 ubiquitin ligase, RAD18, to promote its association with TLS polymerases and subsequent damage bypass (16). TLS polymerase have less well-defined function in other DNA repair pathways that occur independently of PCNA (17–20). HCMV also encodes a viral DNA polymerase processivity factor, *UL44*, which is essential for virus replication (21). Like PCNA, pUL44 is a processivity factor that interacts with pUL54 to maintain polymerase-DNA interactions. Comparing the two, both PCNA and pUL44 are sliding clamps that associate with DNA, but they share no sequence similarity (22). Additionally, PCNA is a homotrimer that undergoes extensive post-translational modifications to regulate interacting partners (23). By contrast, pUL44 is a homodimer with few characterized post-translational modifications and viral interacting partners (24–26). Despite these differences, pUL44 and pUL54 are thought to function similarly to PCNA and replicative cellular B-family polymerases as a clamp-polymerase complex to facilitate vDNA synthesis (21, 27). We sought to better understand the role of PCNA in HCMV replication.

We previously observed that HCMV infection in fibroblasts induces monoubiquitination of PCNA and, in line with observations from others, re-localization of PCNA to viral RCs (14, 28–30). This observation was consistent with the roles we found for TLS polymerases in regulating HCMV genome integrity and replication. However, the significance of PCNA to HCMV genome synthesis, integrity, and replication remains to be defined. Here, we found that PCNA restricted viral replication in the TB40/E strain in a manner that was dependent on PCNA and its modification at the K164 residue. Homing in on this monoubiquitination, we found that accumulation of mUb-PCNA depended on viral DNA synthesis. Further, PCNA and mUb-PCNA localized to distinct subdomains in RCs relative to replication forks and viral proteins important for vDNA synthesis. However, unlike the TLS polymerases that mUb-PCNA would presumably recruit, PCNA and mUb-PCNA were not required for protecting the vDNA from large rearrangements. Instead, we found that PCNA contributed to genome diversity through generating SNVs on viral DNA, similar to Y-family TLS polymerases. These results suggest that, while the SNVs generated on viral DNA by TLS polymerases are PCNA-dependent, TLS polymerase-moderated genome stability occurs independently of PCNA. Altogether this work uncovers specific contributions of PCNA to HCMV infection and highlights the complexity of this virus-host interaction.

## RESULTS

### mUb-PCNA increases with viral DNA synthesis

Given the role of TLS polymerases to HCMV infection and their dependence on mUb-PCNA in host cells, we sought to build on our findings by further characterizing mUb-PCNA in the context of HCMV infection. To better define the relationships between PCNA, mUb-PCNA, and vDNA synthesis, we inhibited viral DNA synthesis with phosphonoacetic acid (PAA) and analyzed the accumulation of PCNA and mUb-PCNA. PAA is a small molecule that binds the viral polymerase, pUL54, and blocks the pyrophosphate binding site, ultimately inhibiting polymerase activity (31, 32). Cells were treated with PAA at the onset of infection (TB40/E-WT or mock). In mock-infected cells mUb-PCNA decreased as cells grew to confluence, and PAA treatment had no effect on this over the 96 hours post infection (hpi) time course (Fig. 1A, quantified in 1B). HCMV-infected cells accumulated mUb-PCNA as infection progressed, consistent with previous findings (14), while levels did not change in PAA-treated cells (Fig 1A, quantified in 1C). Further, unlike mock-infected cells, HCMV infection maintained mUb-PCNA despite increasing cell confluence. These data suggest that HCMV vDNA synthesis drives accumulation of mUb-PCNA.

**Figure 1.**
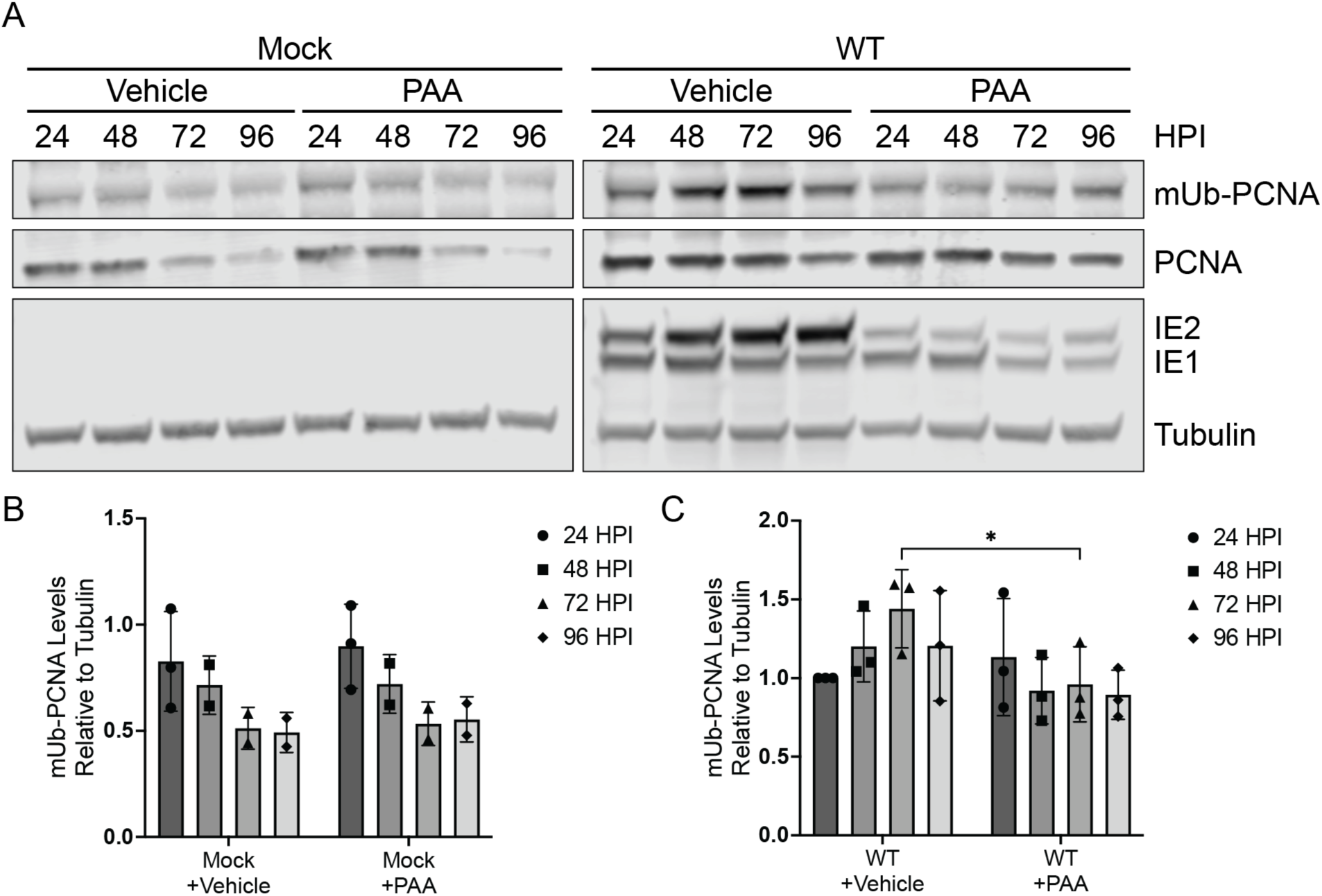
mUb-PCNA increases with viral DNA synthesis. Fibroblasts were mock-infected or infected (MOI = 1) with TB40/E-WT virus over a 96-hour time course. Immunoblotting was performed on whole cell lysates collected at the indicated time points. (A) mUb-PCNA and PCNA were detected using monoclonal antibodies specific to each and secondary antibodies conjugated to DyLight™ 680 (mouse) or 800 (rabbit). IE1/2 are immediate early proteins that serve as a marker for progression of virus replication. Tubulin serves as a loading control. mUb-PCNA levels relative to tubulin were quantified in (B) mock-infected and (C) TB40/E-infected conditions and normalized to the 24 hpi time point. Statistical analysis was performed by two-way ANOVA with Tukey’s multiple comparisons test. Asterisks (*P <0.05) represent statistically significant differences determined in three independent experiments.

### PCNA restricts HCMV TB40/E replication

Given that mUb-PCNA is associated with viral DNA synthesis, we hypothesized that PCNA is functionally important for viral replication. We stably disrupted PCNA expression via shRNA depletion under growth arrest conditions in order to avoid replication stress and cell death due to depletion of this essential host factor. Compared to cells expressing shRNA against firefly Luciferase (Luc, non-targeting control), we achieved ∼70% knockdown of PCNA protein over multiple independent experiments (Fig. 2A). Compared to the Luc control, depletion of PCNA resulted in a ∼2 log increase in virus yield, suggesting that PCNA restricts HCMV replication (Fig. 2B). Consistent with this, we also measured viral genome copy number at 15 days post infection and observed an increase with depletion of PCNA (Fig. 2C). Cells depleted of PCNA infected at a multiplicity of infection (MOI) of 1 and collected over a 96-hour time course exhibited a 4-fold increase in viral titers (Fig. 2D), but no significant increase in viral genomes (Fig. 2E). Therefore, PCNA restricts HCMV replication and genome synthesis, and the PCNA-mediated restriction of viral genome synthesis and progeny production was less apparent at higher MOIs of infection but not fully overcome.

**Figure 2.**
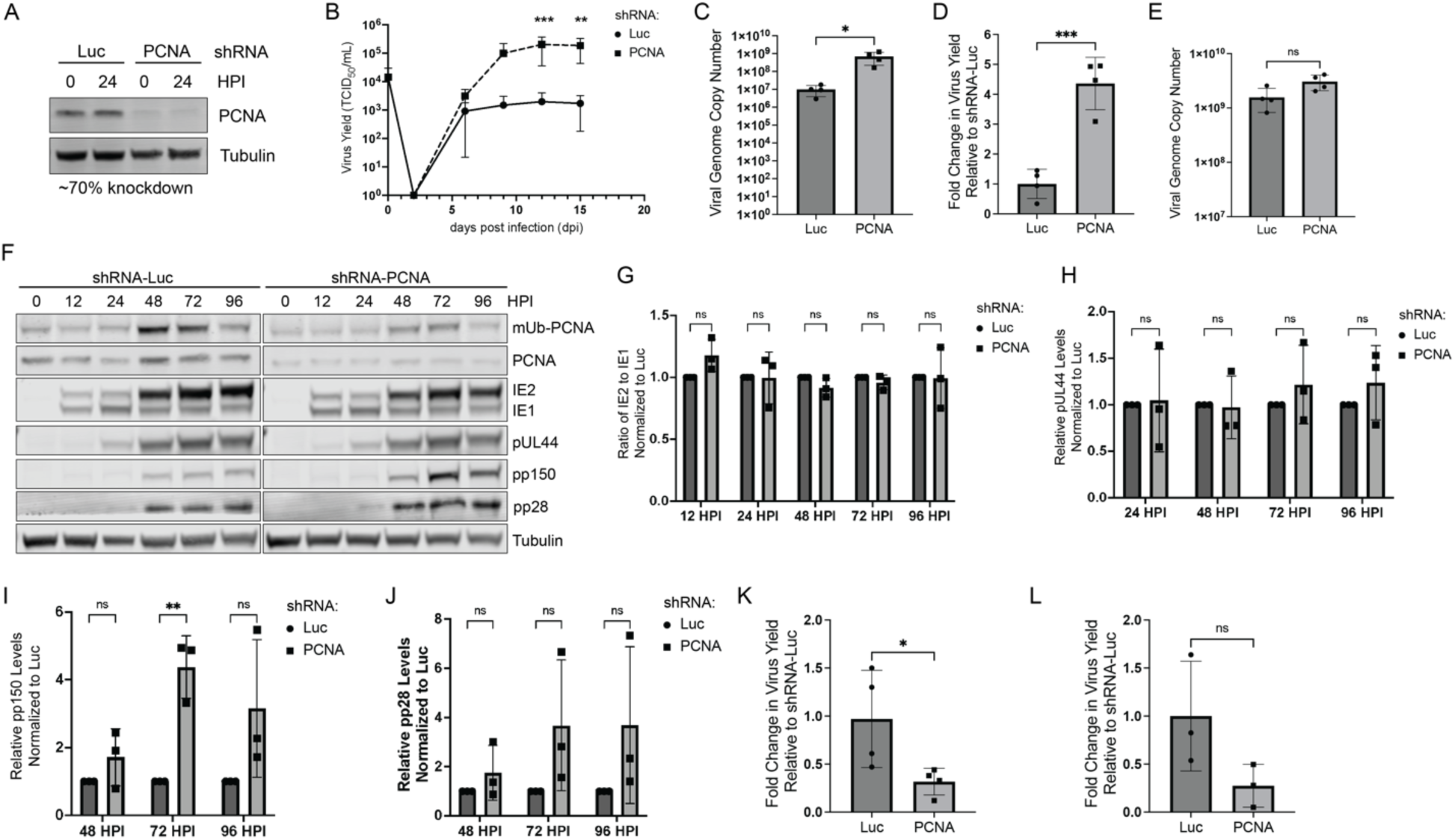
PCNA restricts HCMV TB40/E replication. (A-C) Growth-arrested fibroblasts expressing shRNA against PCNA or Luciferase (Luc, non-targeting control) were infected with TB40/E-WT at an MOI of 0.02. (A) Whole cell lysates were collected at the time of infection (0 hpi) and at 24 hpi and then immunoblotted. To confirm knockdown, PCNA was detected using a monoclonal antibody. An average knockdown of 70% was achieved over multiple independent experiments. (B) Virus yield at 15 dpi were determined by TCID_50_ and normalized relative to the Luc control. (C) Viral genome copy number at 15 dpi was determined by qPCR using a TB40/E BAC standard curve and primer set designed for the region of the viral genome encoding the β2.7 transcript. (D-F) Growth-arrested fibroblasts expressing shRNA against PCNA or Luc were infected with TB40/E-WT at an MOI of 1. (D) Virus yield at 96 hpi was determined by TCID_50_ and normalized relative to the Luc control. (E) Absolute viral genome copy number was determined by qPCR using a standard curve. (F) Whole cell lysates were collected over a 96 hour time course (0 hpi = time of infection) and immunoblotted. To confirm knockdown, PCNA and mUb-PCNA were detected using monoclonal antibodies to each. The following proteins were also detected as markers for the viral gene expression cascade: IE1/2 (immediate early), pUL44 (early), pp150 and pp28 (late) with secondary antibodies conjugated to DyLight™ 680 (mouse) or 800 (rabbit). Tubulin serves as a loading control. (G-J) Quantification of immunoblots in *F* for (G) IE1/2, (H) pUL44, (I) pp150 and (J) pp28 with PCNA knockdown normalized to Luc for each time point. (K) Growth-arrested fibroblasts expressing shRNAs were infected with HSV-1 at an MOI of 0.01 and virus yields were determined relative to the Luc control at 33 hpi. (L) Growth-arrested fibroblasts expressing shRNAs were infected with HCMV AD169-GFP (WT) at an MOI of 1, and virus yield was determined relative to the Luc condition at 96 hpi. For statistical analysis, significance was determined by two-way ANOVA with Tukey’s multiple comparisons test (B), two-way ANOVA with Sidak’s multiple comparisons test (G-J) or an unpaired t test (C-E, K-L). Asterisks (*P <0.05, **P <0.01, ***P <0.001) represent statistically significant differences determined in a minimum of three independent experiments.

To understand how PCNA affects viral gene expression, we further analyzed the impact of PCNA on viral gene expression (Fig. 2F). Comparing between Luc and PCNA knockdown, we observed no differences in immediate early proteins (IE1/2) or a representative early protein, pUL44. Protein levels of IE1/2 and pUL44 from three independent experiments are quantified in Figures 2G and 2H, respectively. Therefore, PCNA does not impact viral gene expression early during infection. However, the late protein, pp150 was increased at 72 hours post infection (hpi), although increases in another late protein, pp28, did not reach statistical significance relative to the Luc control (Fig. 2I-J). Taken together, these data suggest that the restriction imposed by PCNA on vDNA synthesis is reflected in reduced viral gene expression late in infection.

Our observation that PCNA restricts HCMV replication is surprising considering that PCNA is re-localized to HCMV RCs and supports viral DNA synthesis for other herpesviruses (33–35). Specifically, PCNA is required for replication of the alpha-herpesvirus, herpes simplex virus 1 (HSV-1) (35, 36), a finding that we recapitulate when we infect PCNA-depleted cells with HSV-1 (Fig. 2K). Further, PCNA depletion has been shown to decrease genome synthesis in the lab adapted AD169 strain of HCMV (30), which we also observe when measuring virus yield (Fig. 2L). These results suggest that the restriction imposed by PCNA on HCMV replication is due to genes or attributes specific to low-passage strains of HCMV.

### K164 modification on PCNA mediates restriction of HCMV TB40/E replication

To build upon our finding that PCNA restricts HCMV TB40/E replication, we sought to determine the significance of mUb-PCNA during infection. In response to DNA damage, PCNA is monoubiquitinated on lysine 164 (K164) by the E3 ubiquitin ligase, RAD18, facilitating interactions with TLS polymerases (37, 38). PCNA is also modified on K164 by polyubiquitination to activate an alternate, error-free DNA lesion bypass through a post-replicative, template-switching mechanism (37, 39, 40). Further, K164 may also be modified by small ubiquitin-like modification (SUMOylation) to antagonize homologous recombination DNA repair (41, 42). In order to investigate the significance of post-translational modifications on the K164 residue, we generated two shRNA-resistant PCNA constructs through wobble codon mutagenesis: wild-type PCNA and a mutant in which K164 is mutated to arginine (K164R), preventing both ubiquitination and SUMOylation on this residue.

To determine how K164 modification on PCNA impacts HCMV replication, we generated lentivirus particles to deliver these constructs alongside shRNA in an attempt to rescue phenotypes associated with PCNA depletion. While overexpression of wild-type PCNA resulted in monoubiquitination of PCNA, expression of the K164 variant of PCNA resulted in minimal detection of mUb-PCNA as expected (Fig. 3A). Cells expressing these constructs were infected at an MOI of 1 and collected at 96 hpi. For controls, we also compared cells expressing shRNA against Luc or PCNA with no rescue (Empty).

**Figure 3.**
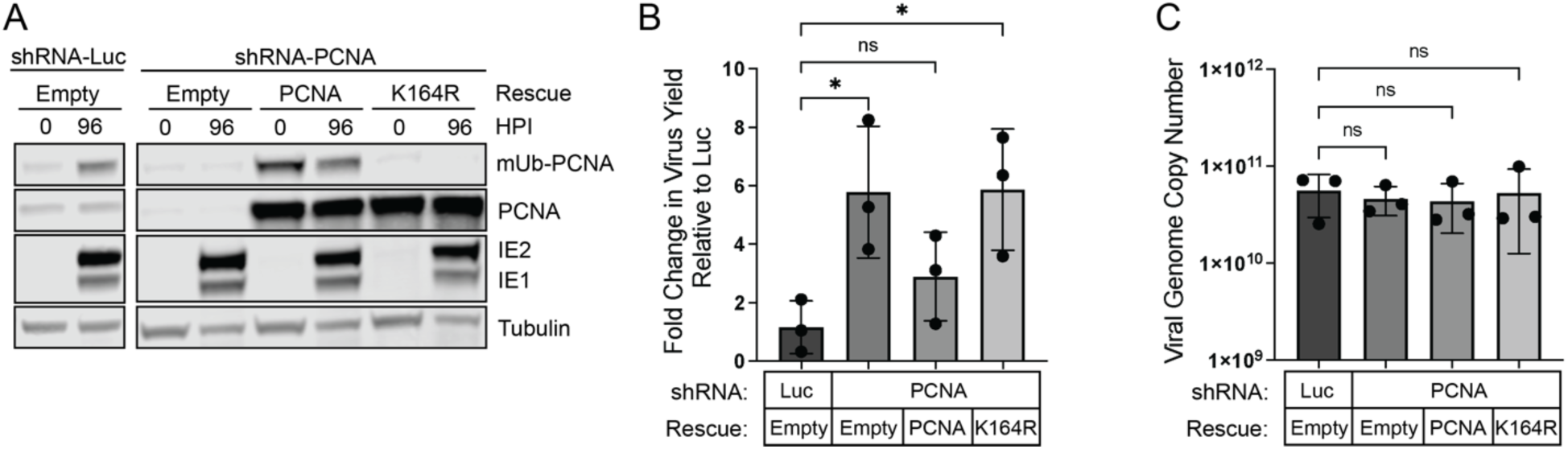
PCNA K164 modification mediates restriction of HCMV replication. Growth-arrested fibroblasts were transduced with lentiviral particles expressing shRNAs (targeting PCNA or Luciferase). Simultaneously, cells were transduced to overexpress shRNA-resistant PCNA (wild-type or mutant containing a lysine-to-arginine mutation at amino acid 164 [K164R]) or an empty overexpression vector control. Cells were infected with HCMV at an MOI of 1 at 48 hours post transduction and collected at 96 hpi. (A) To confirm protein knockdown and rescue, whole cell lysates were collected at the time of infection (0 hpi) or at 96 hpi. mUb-PCNA, PCNA, tubulin, and IE1/2 were detected using monoclonal antibodies as shown with secondary antibodies conjugated to DyLight™ 680 (mouse) or 800 (rabbit). (B) Virus yields were measured by TCID50 and normalized relative to Luc. (C) Viral genome copy number was determined by qPCR using a TB40/E BAC standard curve and primer set designed for the region of the viral genome encoding the b2.7 transcript. Statistical significance was determined by one-way ANOVA with Tukey’s multiple comparisons test. Asterisks (**P <0.01) represent statistically significant differences determined in independent experiments.

As expected, PCNA knockdown with empty rescue recapitulated a ∼five-fold increase in virus yield over Luc control conditions. In comparison, PCNA knockdown with wild-type PCNA expression yielded no significant change in virus replication compared to Luc, demonstrating a partial phenotypic rescue (Fig. 3B). PCNA-K164R rescue resulted in replication at levels similar to PCNA depletion alone, suggesting that PCNA-mediated restriction of HCMV replication depends on modification of K164. Despite the effect on virus replication, PCNA K164R did not impact viral genome copy number (Fig. 3C), similar to our results with PCNA knockdown at this MOI (Fig. 2E). Therefore, PCNA restricts HCMV replication in a manner dependent on modification at the K164 residue, although we cannot differentiate the importance of ubiquitination or SUMOylation of this residue.

### mUb-PCNA localizes to distinct replication compartment subdomains

PCNA localizes to viral replication compartments (14, 28–30), suggesting a role for PCNA in vDNA synthesis. However, the virus encodes its own functional homologue of PCNA, pUL44, that is important for processivity of the viral DNA polymerase, pUL54 (21, 43, 44). To better understand the interplay between pUL44 and either PCNA or mUb-PCNA, we analyzed their localization in viral RCs and colocalization to replication forks. Active sites of vDNA synthesis labeled with EdU have been localized to the periphery of RCs with pUL44 in HCMV infection (45), suggesting that viral DNA synthesis occurs at the periphery. However, others have observed active DNA synthesis throughout HCMV and HSV-1 RCs using either 5-ethynyl-2’-deoxyuridine (EdU) or 5-ethynyl-2’-deoxycytidine (EdC) nucleotide analogs (36, 46, 47). We used this technique to analyze the association of PCNA and mUb-PCNA with sites of active synthesis of vDNA relative to pUL44 or the HCMV single-stranded DNA binding protein, pUL57, which also localizes to RCs (48). To ensure labeling only of viral RCs, fibroblasts were growth arrested prior to infection and maintained in serum-free conditions. At 48 hpi, we pulsed cells with EdU for 10 min and localized PCNA, mUb-PCNA, pUL44, and pUL57 by indirect immunofluorescence to assess their association with EdU-labeled replication forks.

EdU labeling was specifically incorporated throughout the RCs of infected cells (Fig. 4). PCNA exhibited stronger colocalization with EdU than either pUL44 (Fig. 4A) or pUL57 (Fig. 4B). However, mUb-PCNA was more distantly associated with EdU than pUL44 (Fig. 4C) or pUL57 (Fig. 4D), suggesting a possible role post-synthesis. Because we did not observe EdU incorporation at the periphery of RCs as previously reported by Strang et al. (45), we analyzed EdU labeling in cells infected with the AD169 laboratory-adapted strain in case virus strain accounted for these differences. EdU was incorporated throughout the RCs similarly to TB40/E infection (Fig. S1). The differences between the observations reported here and those by Strang et al. may be due to cell type or MOI differences. However, incorporation of EdU throughout RCs has been reported by others in HCMV and HSV-1 infection (36, 46, 47).

**Figure 4.**
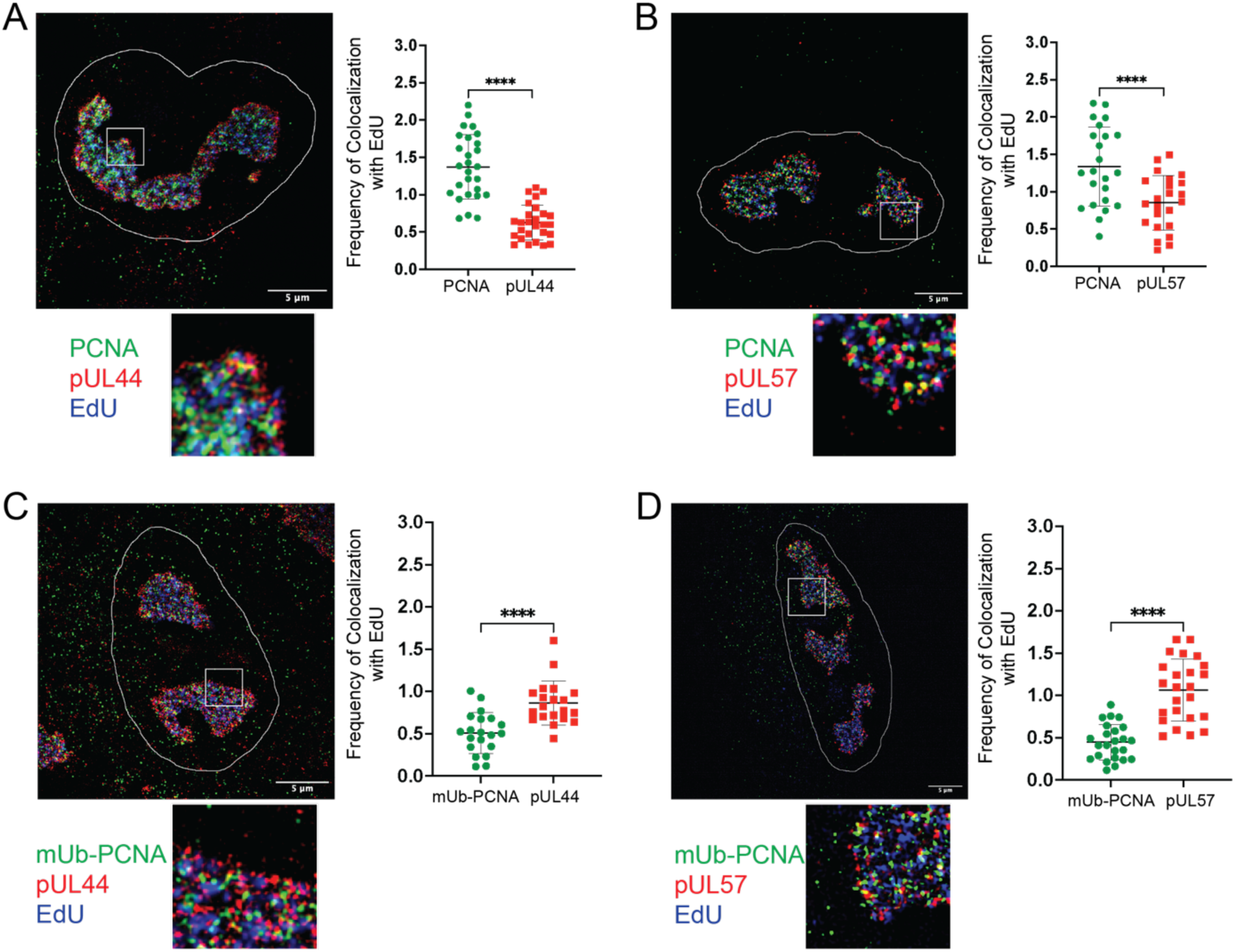
mUb-PCNA localizes to distinct replication compartment subdomains. Fibroblasts were serum-starved and then infected with TB40/E-WT at an MOI of 1. At 48 hpi, cells were pulsed with 10 µM EdU for 10 minutes, CSK-extracted, and fixed. All coverslips were washed and then a click reaction was performed to conjugate EdU and Alexa Fluor 647 (blue) for detection. Indirect immunofluorescence was then carried out using monoclonal antibodies to the indicated proteins: (A) PCNA and pUL44, (B) PCNA and pUL57, (C) mUb-PCNA and pUL44, (D) mUb-PCNA and pUL57 with secondary antibodies conjugated to Alexa Fluor® 546 (green) or 488 (red). DAPI-stained nuclei (not pictured) were outlined using Fiji/ImageJ software. An enlargement of the boxed area is shown below each image. Images were captured using a Zeiss Elyra S.1 super-resolution microscope. Scale bar, 5 µm. The frequency of colocalization between host or viral proteins and EdU was quantified using Nikon NIS Elements software (see Materials and Methods) and is shown to the right of each image with each point representing a cell. Statistical significance was determined by a paired t test. Asterisks (****P <0.0001) represent statistically significant differences determined in three independent experiments.

### PCNA contributes to HCMV genome diversity but not integrity

Our findings thus far point to a role for PCNA in regulating HCMV genome synthesis and replication. As an essential factor for eukaryotic DNA replication and repair, we wondered the extent to which PCNA influences viral genome integrity. In previous studies, we found that TLS polymerases are required for HCMV genome integrity (14). While TLS polymerases can function in DNA repair independently of PCNA (17, 19, 20), PCNA and its mUb is critical to recruit TLS polymerases to lesions for canonical translesion synthesis (15, 23). Building on findings presented here, we sought to determine if depletion of PCNA alone was sufficient to compromise viral genome integrity in HCMV TB40/E infection and if genome integrity depended on modification at K164R. To assess this, we sequenced genomic DNA extracted from HCMV-infected fibroblasts expressing shRNAs against Luciferase or PCNA with expression of an empty vector or PCNA K164R as described (Fig. 3A) and quantitated novel DNA junctions (inversions, duplications, and deletions) and single nucleotide variations (SNVs; point mutations, small deletions, and insertions) arising in synthesized viral genomes compared to the parental virus stock used for infection. Strikingly, we found that neither depletion of PCNA nor rescue with the K164R mutant significantly impacted large genomic rearrangements compared to the Luc control (Fig. 5A, quantified in 5B). This was a surprising result given the role for PCNA in recruiting TLS polymerases. By contrast, depletion of TLS polymerases, increased inversions, duplication and deletions across the viral genome (14), However, similar to depletion of TLS polymerases (14), depletion of PCNA resulted in fewer SNVs on the viral genome, but SNVs are generated independently of modification at K164R (Fig. 5C). Taken together, these results suggest that PCNA is dispensable in the maintenance of HCMV genome integrity but could drive genome diversity.

**Figure 5.**
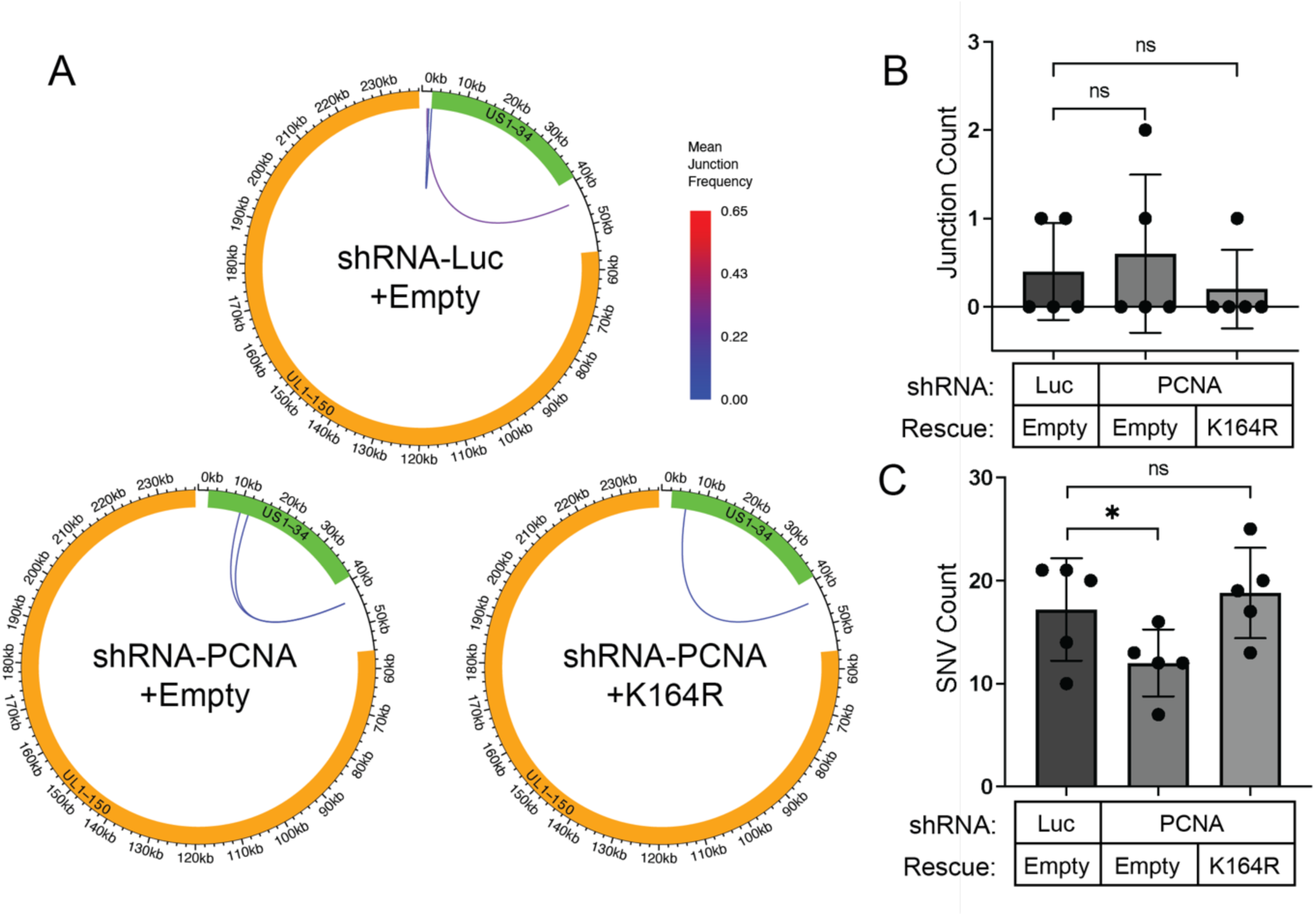
PCNA contributes to HCMV genome diversity but not integrity. Growth-arrested fibroblasts were transduced with lentiviral particles expressing shRNAs against PCNA or Luciferase at an MOI of 3. Simultaneously, cells were transduced to overexpress shRNA-resistant PCNA-K164R or an empty overexpression vector control. Forty-eight hours later, media was refreshed with puromycin at 2 μg/mL; 24 hours later, cells were infected with TB40/E-WT at an MOI of 1 and total DNA was isolated at 96 hpi for sequencing. Sequences from each knockdown condition as well as from the virus stock used for infection were aligned to the TB40/E-GFP reference genome. (A) Mean novel junction frequency within each condition. HCMV genomic coordinates are plotted along the circular axis in graphs for each condition and the UL (orange) and US (green) regions of the genome are marked. The arcs connect novel junction points detected at the average frequency for the given condition indicated by the color scale. (B) Quantification of the number of novel junctions (inversions, deletions, duplications) detected per sample (n = 5) for each condition. (C) Quantification of the number of novel SNVs (point mutations, deletions, or insertions) detected per sample (n = 5) for each condition. Statistical significance was determined by pairwise two-sided exact Poisson tests and adjusted using Bonferroni correction. Asterisks (*P <0.05) represent statistically significant differences determined in independent experiments.

## DISCUSSION

As intracellular parasites, all viruses rely on host cell factors for efficient replication. Even complex DNA viruses, like herpesviruses, hijack their host’s machinery despite encoding similar factors of their own. Our current knowledge of the involvement of host factors in the HCMV replicative program, especially at the step of vDNA synthesis, is incomplete. Building on our previous findings that TLS polymerases are recruited to viral RCs and modulate HCMV replication and genome integrity (14), we set out to define roles of the host DNA processivity factor and polymerase binding partner, PCNA, in HCMV replication. While PCNA is best known for its role in increasing polymerase processivity, it also plays important role in DNA repair (15). Post-translational modifications, such as ubiquitination or SUMOylation, direct its engagement with repair processes. We show that HCMV infection induces monoubiquitination of PCNA, a modification important for recruiting TLS polymerases. PCNA activity on vDNA restricts vDNA synthesis and HCMV replication and post-translational modifications (i.e., monoubiquitination or SUMOylation) at the K164 residue are important for this restriction (Fig. 3B). Despite having a role in viral DNA synthesis, PCNA depletion did not affect viral genome integrity (Fig. 5 A and B), as depletion of TLS polymerases does (14). However, PCNA was required for generating SNVs on viral DNA, similar to Y-family TLS polymerases, ρι, κ, and 1 (Fig. 5C), suggesting that TLS polymerases function on the viral genome by both PCNA-dependent and -independent mechanisms. Additionally, unmodified PCNA and mUb-PCNA were differentially associated with sites of active vDNA synthesis (Fig. 4), further emphasizing the multifaceted role of PCNA in regulating HCMV replication. This work highlights PCNA as a restriction factor to HCMV replication with complex roles that remain to be defined.

With no enzymatic activity, PCNA undergoes multiple post-translational modifications to direct chromatin binding and association with interacting partners. Of these modifications, monoubiquitination by the E3 ubiquitin ligase, RAD18, is important for the recruitment of specialized polymerases to bypass obstructive DNA lesions on cellular DNA (16, 38). Our observation that induction of mUb-PCNA is associated with vDNA synthesis (Fig. 1) suggests this is a virus-driven phenomenon. However, the induction of mUb-PCNA may also be a host response to virus infection. For example, increased virus replication is associated with increased oxidative stress (49, 50), and mUb-PCNA induction has been associated with oxidative stress (51, 52). Additionally, induction of mUb-PCNA and its localization to viral RCs could be a response to replication stress that arises due to vDNA synthesis in the nucleus. In this scenario, the association of PCNA with viral genomes might interfere with vDNA synthesis machinery (i.e., pUL54 and pUL44) and ultimately cause a restriction to vDNA synthesis. To further understand the nature and significance of PCNA-K164, future studies should examine the host and or viral proteins that interact with PCNA and regulate post-translational modifications to direct PCNA function during infection.

We report that PCNA restricts HCMV-TB40/E replication based on observations that shRNA knockdown of PCNA resulted in increased virus yield (Fig. 2). This was a surprising finding considering that HCMV induces PCNA expression (53, 54) and PCNA localizes to RCs (28, 55) with newly synthesized viral DNA (Fig. 5). Further, PCNA facilitates the replication of other herpesviruses, HSV-1 (35) and EBV (56). For HSV-1, PCNA plays a key role by recruiting viral and cellular factors to sites that facilitate viral genome synthesis (36). Various distinctions between HSV-1 and HCMV, including genome size and differences in the viral processivity factors and kinetics of replication, may underlie how these viruses differentially utilize PCNA. Further, our findings with the low-passage TB40/E strain stand in contrast to previous findings showing that PCNA knockdown decreased HCMV viral genome production in studies using the AD169 strain (30). While we observe that PCNA restricts viral genome synthesis in low MOI infections of TB40/E (Fig. 2C), it is possible that strain differences account for the discrepancy in results. In our study, PCNA knockdown in AD169 infection did not produce a significant change is virus replication (Fig. 2L). AD169 notably lacks 15 kb of viral DNA including many genes important for modulating viral replication for establishment of latency (57). Thus, it is conceivable that some of these genes modulate PCNA in a way that restricts virus replication.

For example, pUL145 has been reported to degrade helicase-like transcription factor (HLTF) (58), a multi-functional protein that contains a RING domain which interacts with RAD18 to promote polyubiquitination on PCNA-K164 (40, 59). The strain-dependent differences and the mechanisms by which PCNA restricts HCMV infection remain to be defined.

Multiple studies have demonstrated that regulation of PCNA and its post-translational modifications are important for other herpesvirus infections. During infection by the gamma-herpesvirus, Epstein Barr virus (EBV), PCNA is deubiquitinated by the viral enzyme, BPLF1 (60). Similarly, PCNA ubiquitination is induced during productive HSV-1 infection and antagonized by the viral deubiquitinating enzyme (DUB), UL36USP (61). Both of these studies showed that deubiquitination of PCNA was important for downregulating polρι recruitment to sites of DNA damage outside the context of infection, but the role of PCNA ubiquitination during viral infection remains to be determined. Notably, Whitehurst et al. show that a PIP domain is conserved among herpesvirus-encoded DUBs, suggesting that these viral enzymes could regulate PCNA. HCMV pUL48 has DUB activity (62); however, a role for pUL48 or host DUBs in deubiquitinating PCNA during infection has yet to be defined. It is possible that herpesvirus infections commonly induce mUb-PCNA but its significance to infection varies among different viruses.

PCNA monoubiquitination at K164 is thought to primarily coordinate the DNA damage bypass pathway, TLS. We previously found that TLS polymerases are involved in HCMV infection, whereby Y-family insertion polymerases ρι, κ, and 1 restricted replication, and Rev1 and ζ were required for efficient virus replication (14). Like the insertion TLS polymerases, PCNA restricts HCMV replication, suggesting they could work together through canonical TLS to achieve this effect. Further, similar to previous findings with depletion of TLS polymerases ρι, κ, and 1 (14), SNVs on vDNA were decreased with depletion of PCNA (Fig. 5C). These observations suggest that PCNA-regulated TLS occurs on the HCMV genome to drive SNVs, consistent with the error-prone nucleotide insertion function of these polymerases (17). Surprisingly, however, PCNA knockdown and rescue with the K164R mutant did not impact SNV counts, suggesting that, while PCNA is important for generation of SNVs, it occurs independently of the mUb-PCNA thought to recruit TLS polymerases to DNA.

In contrast to SNVs, PCNA depletion did not increase novel junctions on the genome, indicating no requirement for HCMV genome integrity (Fig. 5A-B), while depletion of TLS polymerases results in increased junctions, primarily inversions (14). Further, rescue of PCNA knockdown with the K164R mutant failed to restore the PCNA-mediated restriction to viral replication (suggesting a requirement for modification at K164, Fig. 3), but had no effect on genome integrity (suggesting the modification on K164 is dispensable for maintaining integrity, Fig. 5). These data suggest that TLS polymerase regulation of HCMV genome integrity occurs independently of PCNA and its monoubiquitination. The fact that depletion of TLS polymerases increased genomic rearrangements, introducing novel junctions on vDNA primarily through sequence inversions, suggests that TLS polymerases are engaging in homology-directed repair, and this occurs independently of PCNA. The function of TLS polymerases in homology-directed recombination are poorly defined (17) and HCMV offers a new tool for defining these roles. For example, polρι and polκ have been implicated in DNA repair at common fragile sites or difficult-to-replicate regions through a mUb-PCNA-independent mechanism (17, 18). polρι can also function in homology-directed repair pathways, such as break induced replication (BIR), without the involvement of PCNA (19, 20). Intriguingly, the PCNA-independent functions of TLS polymerases are associated with recombination, suggesting that HCMV commandeers Y-family polymerases, but not PCNA, for recombination-dependent repair. This is an intriguing avenue for future studies. However, one possible limitation of these interpretations as they apply to the study is that because we only deplete PCNA levels by 70%, it is possible that the remaining protein is sufficient to permit repair in the presence of TLS polymerases. However, given the phenotypes resulting from the depletion of PCNA on vDNA synthesis, late gene expression, replication and the generation of SNVs, we think this is unlikely.

Further, it remains a possibility that in the absence of PCNA, TLS polymerases are recruited by pUL44 or an alternative factor, such as Rev1 (63–65). This would suggest that PCNA and pUL44 compete for interacting partners and provide a possible explanation as to why PCNA restricts HCMV replication. However, our localization studies (Fig. 4), show poor co-localization of PCNA and pUL44 at EdU-labeled replication forks, and PCNA is more closely associated with replication forks than either pUL44 or pUL57. Future studies will address the roles of other factors, such as viral pUL44 or host Rev1, in TLS polymerase repair activity during infection.

mUb-PCNA is less closely associated with EdU-labeled replication forks than pUL44 or pUL57, suggesting possible post-synthesis roles that are likely not related to TLS (Fig. 4). The differential subnuclear localization of unmodified and mUb-PCNA implies specific, separable functions of PCNA. It is possible that mUb-PCNA is induced to prevent K164 modification by SUMOylation, which suppresses DNA repair by the homologous recombination pathway (41, 42). This would support data in Figure 3 showing that PCNA’s suppressive effect on HCMV TB40/E replication is mediated by K164 modification. However, further investigation is required to attribute this to monoubiquitylation, polyubiquitylation, SUMOylation, or a combination of these modifications. Additionally, ubiquitination of PCNA is linked to its retention on chromatin and regulation of nucleosome deposition (66), a function required for HSV-1 infection (35). As PCNA interacts with a wide array of repair proteins, further investigation is required to define mUb-PCNA-associated host and/or viral proteins to elucidate this function.

In summary, our findings underscore the intricacy of HCMV interactions with its host, especially with respect to DNA synthesis and repair. Through studying the role of PCNA in HCMV infection, we uncovered multiple, distinct functions of PCNA and, by extension, TLS polymerases on viral genomes that deviate from their canonical function in TLS in host cell biology (Fig. 6). While TLS polymerases engage in canonical TLS repair of vDNA, which is likely PCNA-dependent, but mUb-PCNA-independent, they also likely function in a PCNA-independent manner to protect viral DNA from faulty recombination. Additional work is required to understand specific mechanisms of host machinery-mediated repair of viral genomes by TLS polymerases and the role of PCNA. HCMV infects a large diversity of cell types and has a large, complex exogenous genome, which is readily manipulated. Further, the vDNA is synthesized in the context of cell cycle arrest and inhibition of host DNA synthesis. This allows for the knockdown of critical host factors important to synthesis or repair that would otherwise result in stress or cell death, confounding any results for the requirement in host cell or infection biology. Therefore, HCMV offers an exciting tool to further mechanistically define and separate complex, intermingled DNA synthesis and repair pathways.

**Figure 6.**
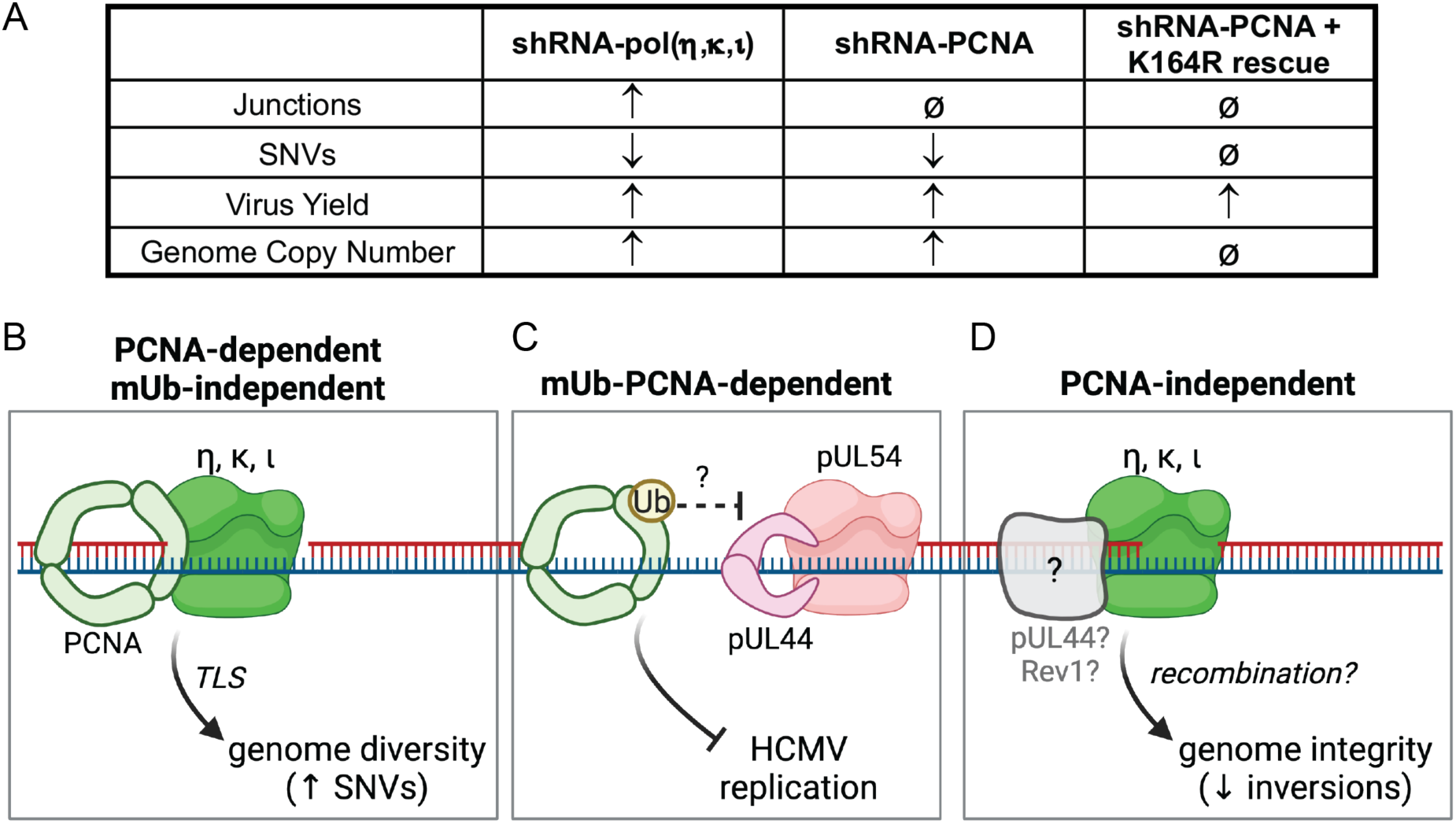
Model for the varied roles of PCNA and TLS polymerases in HCMV DNA synthesis and genome integrity. (A) Summary of observed virus infection phenotypes with host factor knockdown. Table depicts an observed increase (τ), decrease (,), or no change (ø) for each unique condition relative to control (shRNA-Luc). (B) PCNA and TLS polymerases are required for viral genome diversity. In a mUb-PCNA-independent manner, PCNA and error-prone Y-family TLS polymerases ρι, κ, and 1, engage in TLS repair on vDNA, contributing to generation of SNVs and genome diversity. (C) PCNA restricts HCMV replication dependent on its modification at K164. While an exact mechanism remains to be defined, PCNA could compete with or inhibit functions of the viral processivity factor, pUL44, and DNA polymerase, pUL54. (D) TLS polymerases have PCNA-independent functions in HCMV genome integrity. Y-family TLS polymerases ρι, κ, and 1 maintain HCMV genome integrity by preventing inversions on vDNA, likely through recombination-dependent repair. This repair is independent of PCNA and possibly involves an alternative factor, such as pUL44 or Rev1.

## MATERIALS AND METHODS

### Cells and Viruses

Primary human lung MRC-5 fibroblasts (ATCC CCL-171) were maintained in DMEM containing 10% FBS as previously described. Cells were infected with the low-passage HCMV strain, TB40/E, a gift from Dr. Christian Sinzger, which was engineered to express green fluorescent protein from the SV40 promoter.

### RNAi

Control shRNA targeting Luciferase was purchased from Sigma-Aldrich (#SHC007). The shRNA targeting PCNA was constructed in the pLKO.1 backbone. In brief, a 21-mer oligonucleotide (GAATGAACCAGTTCAACTAAC) was generated for the target gene and then cloned into pLKO.1 vector via annealing.

### Lentivirus and transduction

Lentiviral particles for shRNA delivery were cultured in HEK 293T cells as previously described. For transduction, MRC-5 fibroblasts were grown to confluence and growth-arrested by contact inhibition to avoid deleterious effects of PCNA depletion in cells undergoing division. Cells were transduced with lentivirus shRNA or rescue constructs at an MOI of 3 in media containing 1 µg/mL polybrene. At two days post transduction, cells were washed in PBS and then given fresh growth media. At four days post transduction, cells were infected with HCMV as described.

### Plasmids

The genetic sequence for human PCNA was amplified from pME-GFP-PCNA (Addgene #105977). Primers containing a sense mutation for shRNA target sequence were used to amplify shRNA-resistant PCNA. Primers containing K164R substitution were used on this sequence to amplify shRNA-resistant PCNA-K164R. PCNA and PCNA-K164R were then expressed from a pCIG vector. All plasmid sequences were validated by Sanger sequencing. **EdU pulse labeling.** Fibroblasts were seeded onto coverslips in 24-well plates in DMEM containing 10% FBS. The next day, cells were washed three times in PBS and growth medium was replaced with serum-free (0% FBS) culture medium. On the following day, cells were infected with HCMV TB40/E at an MOI of 1. At 48 hours post-infection, half of the media was replaced with serum-free DMEM containing 20 µM EdU for a final concentration of 10 µM EdU. Cells were then incubated at 37°C with 5% CO_2_. After 10 minutes, the coverslips were washed in PBS and then incubated in cold cytoskeletal (CSK) extraction buffer for two minutes (67).

Following CSK extraction, cells were fixed with 100% methanol for 10 minutes at −20°C. After fixation, cells were washed twice with PBS containing 3% BSA. For detection of EdU, the click reaction was performed according to manufacturer instructions (Invitrogen #C10340). Indirect immunofluorescence was subsequently performed as described below.

### Immunofluorescence and super-resolution microscopy

Fibroblasts were seeded onto 12-mm 1.5H high precision coverslips (Marienfeld Superior) in 24-well plates. After EdU pulse labeling (described above), coverslips were processed for indirect immunofluorescence as previously described (14). Briefly, proteins of interest were detected using specific primary antibodies for one hour at room temperature or overnight at 4°C as described in Table 1.

**Table 1.** Antibodies used in this study.

| Antibody | Species | Source | Concentration |
| --- | --- | --- | --- |
| PCNA | Mouse | Santa Cruz, sc-56 | IB 1:1000 |
| PCNA | Rabbit | Cell Signaling Technology (CST), #13110 | IF 1:400 |
| mUb-PCNA | Rabbit | CST, #13439 | IB 1:1000 |
|  |  |  | IF 1:100 |
| α-Tubulin | Mouse | Sigma-Aldrich, #T9026 | IB 1:2000 |
| IE1/2 | Mouse | Thomas Shenk, PhD<br>(Princeton University) | IB 1:100 |
| pUL44 | Mouse | Virusys, #CA006 | IB, IF 1:12,000 |
| pUL57 | Mouse | Virusys, #P1209 | IF 1:100 |
| pp150 | Mouse | William J. Britt, MD (University<br>of Alabama at Birmingham) | IB 1:15 |
| pp28 | Mouse | William J. Britt, MD (University<br>of Alabama at Birmingham) | IB 1:15 |
| Mouse IgG (H+L)<br>secondary, DyLight<br>680 | Goat | Invitrogen, #35519 | IB: 1:6000 |
| Rabbit IgG (H+L)<br>secondary, DyLight<br>800 | Goat | Invitrogen, #SA5-10036 | IB: 1:6000 |
| Mouse IgG (H+L)<br>secondary, Alexa<br>Fluor 488 | Goat | Invitrogen, #A-11029 | IF: 1:3000 |
| Rabbit IgG (H+L)<br>secondary, Alexa<br>Fluor 546 | Goat | Invitrogen, #A-11035 | IF 1:3000 |
| Mouse IgG (H+L)<br>secondary, Alexa<br>Fluor 647 | Goat | Invitrogen, #A-21236 | IF: 1:3000 |

Coverslips were washed 3 times in PBS + 0.05% Tween20 and then incubated in secondary antibodies (AlexaFluor 546, AlexaFluor 647, or AlexaFluor 488 goat anti-mouse or goat anti-rabbit [Invitrogen]) for 30 minutes at room temperature. Coverslips were incubated in DAPI for 5 minutes and then washed three times in PBS + 0.05% Tween20. Finally, coverslips were mounted onto microscope slides (Fisher Scientific) using Prolong Gold Antifade Mounting Reagent (Invitrogen). Images were obtained using a Zeiss Elyra S.1 super-resolution microscope using a Zeiss 63x Plan-Apochromat 1.40NA objective with structured illumination (SR-SIM) processing. Representative single plane images were adjusted for brightness and contrast using Fiji/ImageJ software.

### Colocalization analysis of host and viral proteins to EdU staining in host cell nuclei

Nikon NIS Elements AR 5.42.03 software with the General Analysis 3 (GA3) module was used for image processing and analysis. The following image processing tools were used to achieve accurate segmentation on the EdU, host protein, and viral protein foci channels: Rolling ball background subtraction (radius = 15px) and Median filter (kernel size = 1px). The Bright Spots Detection tool (diameter = 2px) was used to threshold the EdU, Host, and Viral Protein foci. For DAPI, a Gaussian filter (sigma = 8), rolling ball background subtraction (radius = 220px), and gamma correction (gamma = 0.8) were applied. A Signal Intensity threshold was used to threshold host nuclei and the resulting nuclei binary objects were then modified using Grow Regions and Smooth tools to further refine nuclear segmentation. The AND binary operation was performed between DAPI and corresponding EdU, host protein, and viral protein foci to restrict analysis to only the foci present inside each cell nucleus. To quantify the frequency of colocalization between EdU/Host Foci and EdU/Viral foci, respectively, the AND binary operation was performed between these two paired groups and the Object Count function was used to quantify each group.

### Immunoblotting

Whole cell lysates were extracted using RIPA lysis buffer (Pierce) and manual scraping. 50 µg of lysate was loaded onto precast 4-12% bis-tris gels (ExpressPlus™; GenScript) and then proteins were separated by electrophoresis and transferred onto 0.45 µm pore size PVDF membranes (Immobilon®-FL; Millipore). Proteins of interest were detected using primary antibodies described in Table 1 and fluorophore-conjugated secondary antibodies. Images were obtained using a Li-Cor Odyssey CLx scanner and protein levels were quantitated using Image Studio Lite software.

### Genomic sequencing and computational analysis

Fibroblasts were seeded onto 6-cm dishes and transduced with lentiviral constructs as described above. Five technical replicates were seeded for each condition. At four days post transduction, cells were infected with HCMV-TB40/E at an MOI of 1. Virus inoculum was removed at 2 hpi and cells were provided fresh media. At 96 hpi, when maximal cytopathic effect (CPE) was observed, cells were washed with PBS and collected in DNA lysis buffer containing 200 µg/mL proteinase K by manual scraping. After a two hour 55°C incubation for proteinase K digestion, cellular and viral DNA were isolated using phenol-chloroform extraction. DNA was similarly extracted from the virus stock used for infection. All purified DNA was submitted to SeqCenter (Pittsburgh, PA).

Each sample (including virus stocks) was sequenced on a NextSeq 2000 Illumina short read sequencer to yield paired-end short reads. The sequencing reads were aligned to the human reference genome GRCh38 (68) using Bowtie2 v2.5.1 (69) with the following parameters:--very-sensitive for sensitive alignment,--seed 1 for seeding alignment. After alignment, reads aligned to the human genome were filtered out using Samtools v1.17 (70). The following parameters were used:-f UNMAP,MUNMAP to extract reads that were not aligned to the human genome, and -bh to output the filtered alignments in BAM format. HCMV junctions and SNVs were then detected using a two-pass analysis to ensure comparable sequencing coverage between the samples. Non-human reads were first aligned to the reference HCMV genome with breseq v0.38.1 (71) using polymorphism-prediction mode. Using the mean sequencing coverage of reads aligned to the reference from the first breseq run output, the sequencing reads from each sample were subsampled with seqtk v1.4q (https://github.com/lh3/seqtk) using parameter-s 100 and the appropriate respective proportions to yield the same mean coverage equal to that of the sample with the lowest coverage (shRNA-Luc). The subsampled FASTQ files were then used to detect the junctions and SNVs by running breseq in the--polymorphism-prediction mode again. Subsequent data analyses were conducted in R version 4.3.2 (72). The novel junctions and SNVs were obtained by removing any of those found in the virus stock samples from the total junctions and SNVs detected in each experimental sample. A Poisson two-sided test was used to compare junction frequency across experimental conditions, with a junction frequency cutoff of 0.025 employed to refine the selection of relevant sequences. Additionally, circle plots were constructed using ‘circlize’ (version 0.4.16) (73) to visualize the locations and relationships of these sequences, providing a comprehensive view of their distribution and interaction within the genome. This multifaceted approach allowed for a nuanced exploration of genomic junctions, highlighting significant variations and patterns across experimental conditions.

## Data availability

Alignments used for Figure 5 are available at the University of Arizona Research Data Repository (doi: 10.25422/azu.data.27948387). Raw sequence reads have been deposited in the Sequence Read Archive, https://www.ncbi.nlm.nih.gov/sra (BioProject ID: SUB14917988).

## ACKNOWLEDGMENTS

We are grateful to Dr. Jill Dembowski and Dr. Jessica Packard (Duquesne University), Dr. Donald Coen (Harvard University), Dr. Blair Strang (University College of London), Dr. James Alwine (University of Pennsylvania), and Dr. Lynn Enquist (Princeton University) for critical discussion. We acknowledge Emily Jamboretz for technical assistance. The Zeiss Elyra S.1 microscope is part of the Imaging Cores – Optical, which is overseen by the University of Arizona’s Office for Research, Innovation, and Impact (purchase of this instrument was supported by NIH S10 OD019948). We acknowledge the assistance of Douglas Cromey, MS, Co-Manager of the Imaging Cores – Optical. We are grateful for the support of the Nikon Center of Excellence and the Cancer Center Support Grant P30 CA023074 awarded to University of Arizona. This work was funded by grants from the National Institutes of Allergy and Infectious Diseases to FG (AI079059 and AI079059-14S1) and to FG and GB (AI177392 and AI177392-02S1).

