## Supplementary material for "Complex roles for proliferating cell nuclear antigen in restricting human cytomegalovirus replication": Figure S1

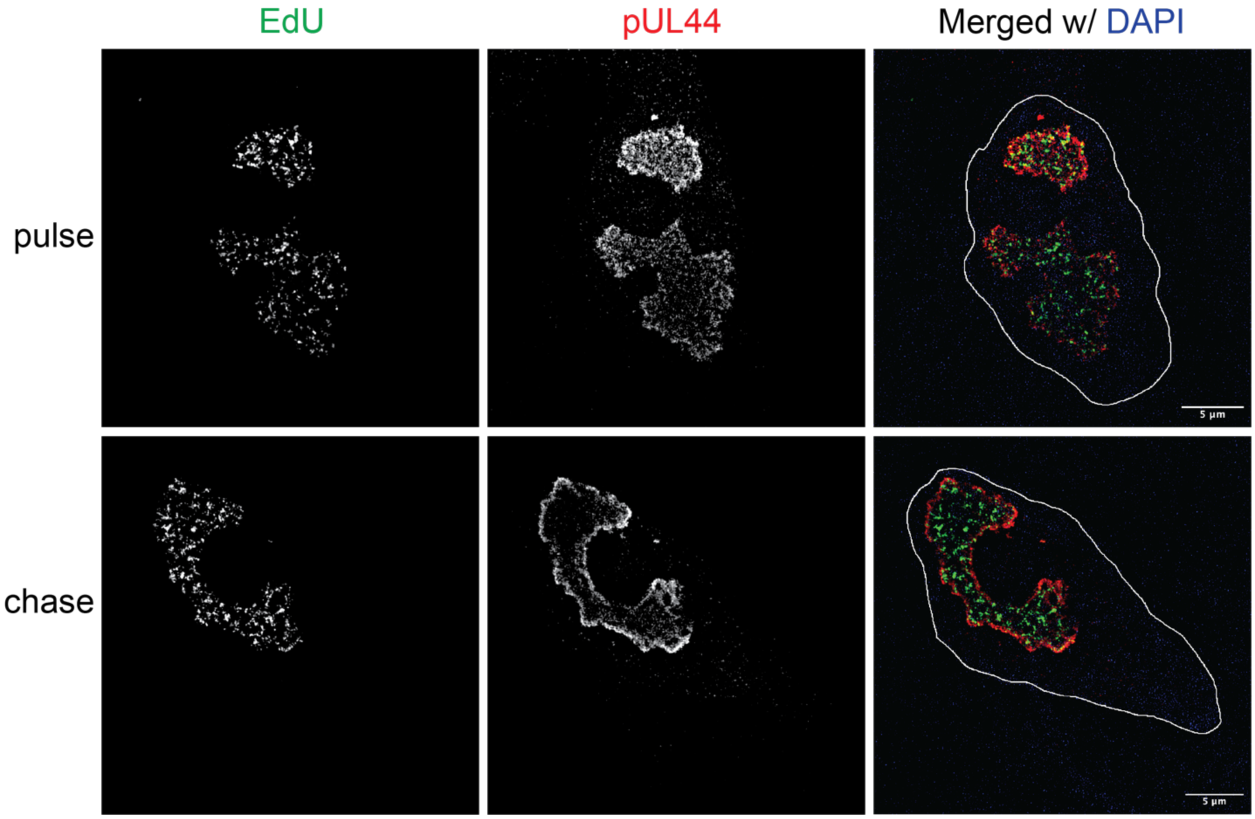


**Figure S1. EdU pulse-chase labeling of viral replication compartments in HCMV AD169 infection.**

Fibroblasts were serum-starved and then infected with AD169-WT at an MOI of 1. At 48 hpi, half of the cells were pulsed with 10 μM EdU for 10 minutes, CSK-extracted, and fixed. Following EdU pulse, the other half of the cells were incubated with 200 μM thymidine for one hour and subsequently CSK-extracted and fixed. All coverslips were washed and then a click reaction was performed to conjugate EdU to Alexa Fluor 647 (shown in green) for detection. Indirect immunofluorescence was then carried out using monoclonal antibodies to pUL44 for detection of viral RCs with a secondary antibody conjugated to Alexa Fluor® 488 (shown in red). DAPI-stained nuclei (individual images not shown) were outlined using Fiji/ImageJ software. Images were obtained using a Zeiss Elyra S.1 super-resolution microscope. Scale bar, 5 μm.
